# Assessing the relationship between neural entrainment and altered states of consciousness induced by electronic music

**DOI:** 10.1101/2024.01.16.575849

**Authors:** Raquel Aparicio-Terrés, Samantha López-Mochales, Margarita Díaz-Andreu, Carles Escera

## Abstract

In electronic music events, the driving four-on-the-floor music appears pivotal for inducing altered states of consciousness (ASCs). While various physiological mechanisms link repetitive auditory stimuli to ASCs, entrainment—a brainwave synchronization through periodic external stimuli— has garnered primary focus. However, there are no studies systematically exploring the relationship between entrainment and ASCs. In the present study, we depart from the finding that entrainment to auditory stimuli peaks for stimulation rates around 2 Hz compared to others. Twenty participants listened to six one-minute electronic music excerpts at different tempos (1.65 Hz, 2.25 Hz, and 2.85 Hz). For each excerpt, they performed cognitive tasks and reported phenomenological experiences related to ASCs through questionnaires. Brain activity was recorded with electroencephalography to assess whether a modulation in entrainment by the beat of electronic music affected objective and subjective proxies of ASCs. Our results revealed a tempo-driven entrainment modulation, peaking at 1.65 Hz. Similarly, participants’ experience of unity during listening to the music was higher for the excerpts at 1.65 Hz, yet no relationship with entrainment was found. Critically, a correlation was found between entrainment and participants’ reaction time. Further studies are granted to explore how individual traits, such as musical training, modulate the relationship.

## Introduction

Every summer since 2005, thousands of people from around the world gather together in Bloom (Flanders, Belgium) to join the Tomorrowland festival. Attendees to this type of electronic dance music events (EDEMs) share the purpose of engaging in transformational experiences in the context of a diversity-friendly environment^[1,2]^. The dynamics of this type of events seem to facilitate their goal^[1]^: the dancing, drumming-like music, sleep deprivation and drug consumption (i.e., the 4D’s) that characterize EDMEs are key elements to produce altered states of consciousness (ASCs)^[3]^. ASCs might enhance the feelings of connectedness, community, and sacrality that EDMEs’ attendees crave^[2]^. However, the physiological mechanisms by which some of the 4D’s facilitate a transition into an ASC remain unknown, being most of the research focused on recreational drugs, especially psychedelics^[4]^. In view of this research gap, we here aim to shed light on the underlying cerebral mechanisms of the 4D’s “drumming” to produce ASCs.

The sonic environment that characterizes EDMEs is reminiscent of that present during shamanic practices^[5]^. During shamanic rituals, intense drumming usually plays during the whole duration of these events to facilitate a shamanic trance state^[6–9]^, which practitioners commonly describe as going into spiritual journeys^[10]^. From a psychological standpoint, trance has been characterized by a narrowed awareness of the immediate surroundings and/or a selective focus on environmental stimuli^[11]^. In modern EDMEs, the disc-jockey takes a similar guiding role to the shaman^[12]^ and the drumming is replaced by electronic music, with an equal strong beat and repeating musical structure that is supposed to entrance the listener^[13]^.

Using repetitive stimulation to facilitate ASCs has not only been described in shamanic rituals and EDMEs, but also in other different cultures and historical periods^[14,15]^. This cross-cultural and cross-temporal commonality suggests the existence of a biological basis that can explain how being exposed to repetitive stimuli might facilitate ASCs^[16]^. Although many explanatory mechanisms have been proposed, neural entrainment is avowed as the most plausible one^[17]^, as it has been shown that brain oscillations eventually match or entrain the phase of external driving stimuli in many sensory domains^[18]^. This match seems to be consequence of an active brain mechanism to ensure a fine processing of the stimuli^[19,20]^, thus engaging the brain’s endogenous activity, that is key to neural processing in a behaviorally relevant manner^[21]^. Additionally, research in beat perception using auditory repetitive stimulation has shown that this brain-stimulus coupling is not only limited to the auditory system, but spreads to the motor system and beyond^[22,23]^. Based on these findings, the drumming in shamanic rituals, or the salient and repetitive beat of electronic music in EDMEs, might entrain several brain regions to frequencies that are not optimal for some cognitive operations. At the behavioral level, such a physiological configuration would translate into a temporary ASC. However, this brain-state of the mind relationship has never been experimentally measured. In this study, we aim to address this previously unaccounted association.

The neuroscientific research in the field of ASCs, specifically that exploring trance, frequently lack measurements of the behavioral correlates associated with these states^[24–27]^, leading to descriptive physiological data. These studies, although being pioneers in the field, did not employ existing tools to explore the invariant phenomenological dimensions of ASCs, such as the 11D-ASC questionnaire ^[28]^. Moreover, the cognitive aspects of ASCs can be measured with traditional cognitive tasks used in psychological research.

In view of these findings, the present study will directly tackle for the first time the assumed relationship between neural entrainment and ASCs. To this aim, we draw upon the finding that neural entrainment to drum sounds and clicks reaches a peak at around 2 Hz compared to other stimulation rates^[29]^. Noteworthy, this physiological property of the brain has never been used to explore how the magnitude of entrainment affects different aspects of human functioning and experience. Specifically, we will measure (1) participants’ entrainment to the beat of naturalistic electronic music at different tempos with electroencephalography (EEG), (2) objective measures of cognitive function, namely executive function and absorption to the music at different tempos, and (3) participants’ subjective experience during the music listening with three subscales of the 11D-ASC. The objective and subjective measures will be employed as proxies to ASCs.

## Methods

The study was approved by the Bioethics Committee of the University of Barcelona and all provisions of the Declaration of Helsinki were followed.

### Participants

A total of 20 naïve, healthy volunteers were recruited from online advertisements at the University of Barcelona (Spain). All participants provided written informed consent and were paid for their participation. The general inclusion criteria comprised no history of auditory, neurological or psychiatric disorders and age between 18 and 35 years. One participant was excluded for being unable to comply with the task instructions, resulting in a final sample of 19 participants of ages ranging from 18 to 22 years (2 male and 17 female, M_age_ = 20). In order to control for previous musical training, participants filled a standard questionnaire we use in our research designed to ascertain both the presence and duration of formal musical training. Out of the 19 participants, six individuals reported prior formal musical education, with a mean duration of 5.833 years (SD = 5.269).

### Stimuli

One-minute-long extracts from “Endless Horizons” by Dhamika (song 1), “El despertar de Joel” by Lab’s Cloud (song 2), “Mind Expander” by Audiomatic (song 3), “Audioslave” by Vertex (song 4), “Check this shit out” by AB (song 5), and “Adagio” by K Complex – Rave Remix (song 6) were used (see supplementary materials). These extracts were carefully selected to include none to minimal vocals and none to minimal beat drops (see Fig. 3a). Each song was modified with v.3.0.0 of Audacity® recording and editing software to include a fade-in and fade-out effect on the first and last seconds, respectively. The approximate tempos of the six songs were determined in a semi-automatic approach (see Section 2.4 for more details). The tempos were: 1.65 Hz or 99 bpm (song 1 and 2), 2.25 Hz or 135 bpm (song 3 and 4), and 2.85 Hz or 171 bpm (song 5 and 6). The six songs could therefore be classified into three tempos: 1.65 Hz, 2.25 Hz and 2.85 Hz. The sound intensity levels were intentionally not equalized across the song extracts because doing so would not effectively address the natural differences in the volume of the beat among the different songs. However, all the extracts were played at the same intensity level.

**Figure 3.**
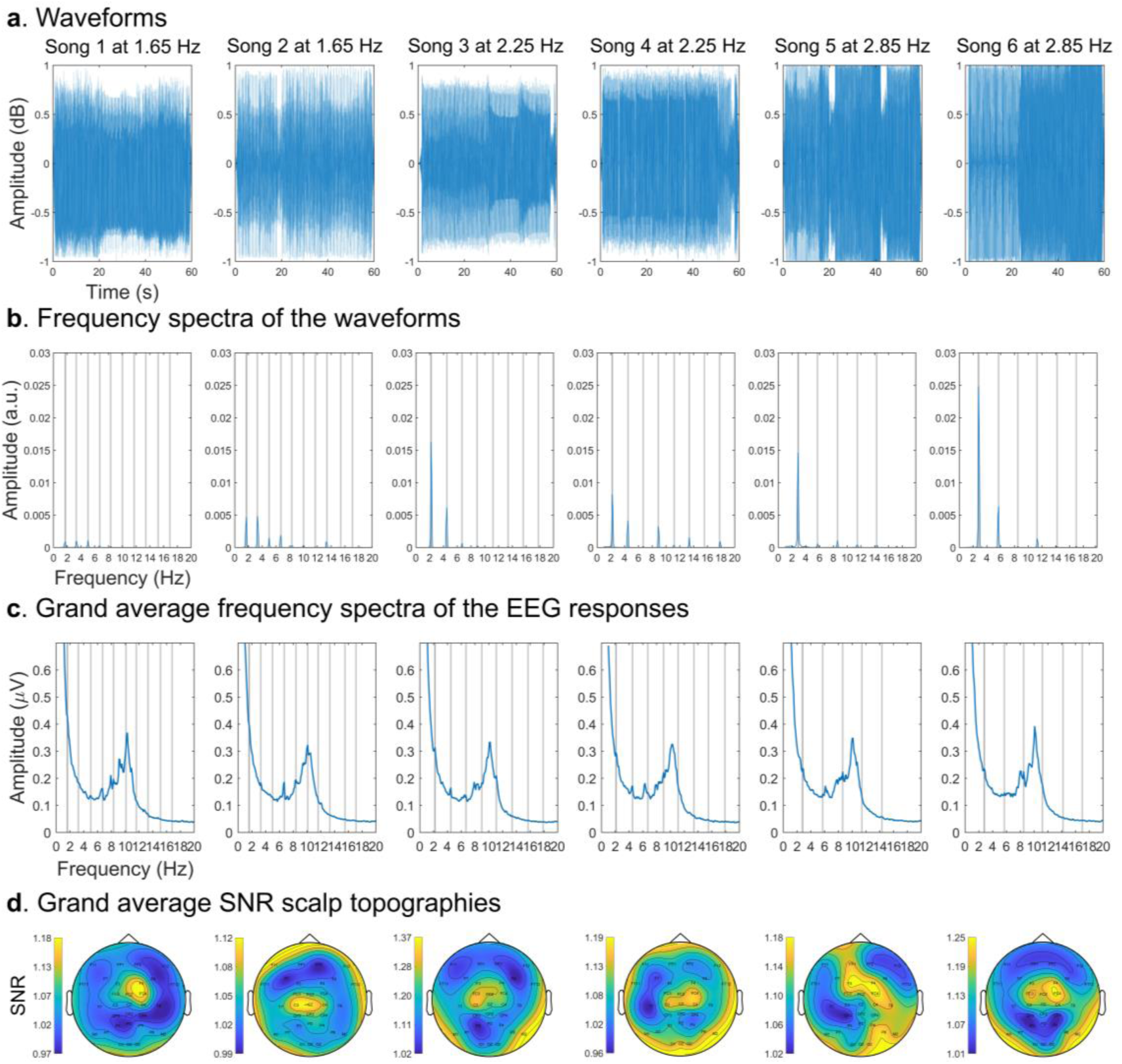
Sound patterns and frequency and spatial representations of the neural responses. (**a**) Waveforms of each song. (**b**) Spectral amplitude of the waveforms. (c) Spectral amplitude of the grand average EEG responses to each song. Gray vertical lines represent the beat frequency and harmonics lower than 20 Hz. (**d**) Scalp topographies of the grand average multi-harmonic SNRs.

### Procedure

Participants sat comfortably in a soundproof Faraday chamber with a screen in front of them, headphones on, a mouse in their dominant hand, and a touch sensitive tablet on their legs. A graphical representation of the sequence of an example trial can be seen in Fig. 1. At the beginning of each trial, participants were presented with a 0.5-second-long pure tone indicating the start of a 10 s silence period. Immediately after, one of the six one-minute-long songs was played at 80 dBs. In total, there were three trials for each song randomized across three blocks for each participant. When the music stopped, participants had to left-click on the mouse as soon as they realized the music was over. After 10 s, a 0.5-second-long pure tone warned participants that the go/no-go task was about to start. Then, the screen displayed a countdown starting from three and framed by a visual shape that could either be a square or a diamond shape. Once the countdown reached one, 150 shapes (either a square or a diamond) were presented sequentially in the screen, following the same timing constraints as in a previous study^[30]^. Specifically, prior to the presentation of each shape, a fixation cross was presented for 200 ms + 0-300 ms jitter. Following the fixation cross, each shape was displayed up to 700 ms or until response. If the shape in the screen matched the one presented during the countdown, participants had to left-click on the mouse (go stimuli). If not, they had to refer from clicking (no-go stimuli). The shape of the go-stimuli was randomized for each participant. The probability of a go-stimuli appearing was fixed in each trial to 75 % to maximize the number of false alarms (i.e., responding when a no-go stimulus is presented)^[31]^. When the go/no-go task was over, participants used the touch-sensitive tablet to fill from 0 to 10 the items presented in a spider chart of three subscales of the 11D-ASC^[28]^, namely Experience of unity, Spiritual experience, and Disembodiment. Following a similar procedure to a previous study^[32]^, the ratings were obtained using three customable spider charts, one for each subscale, with as many radii as items (Fig. 2). A fixation cross in the center of the screen was present during the whole duration of the trials. Previously to the start of the experiment, all participants conducted three practice trials, in which generic elevator music was used as stimuli, to get familiarized with the tasks. EEG was recorded during the whole duration of the trials.

**Figure 1.**
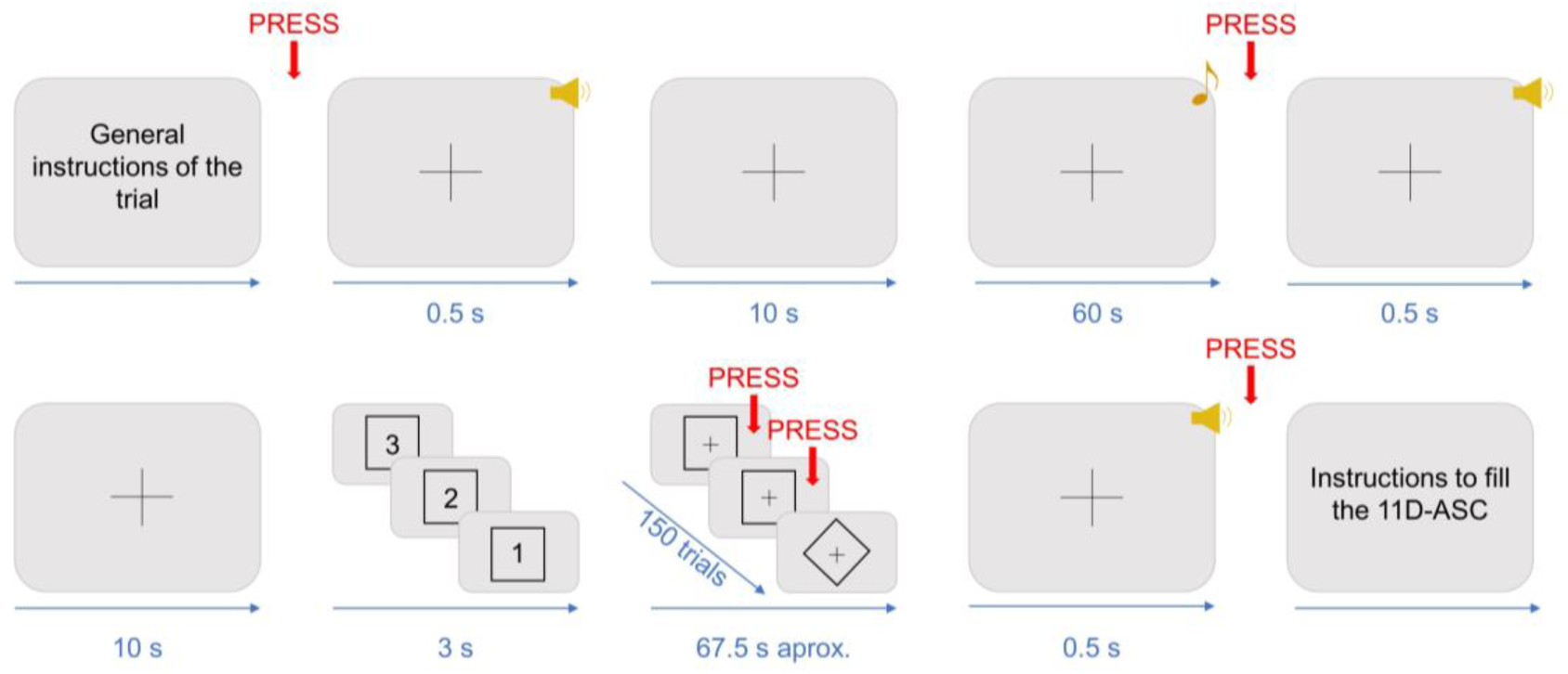
Schematic representation of the experimental design. The figure shows the sequence of motor responses (in red), auditory stimuli (in yellow), and visual stimuli (in black) that participants produced or were presented with during each trial. The timing constraints (in blue) are represented in seconds.

**Figure 2.**
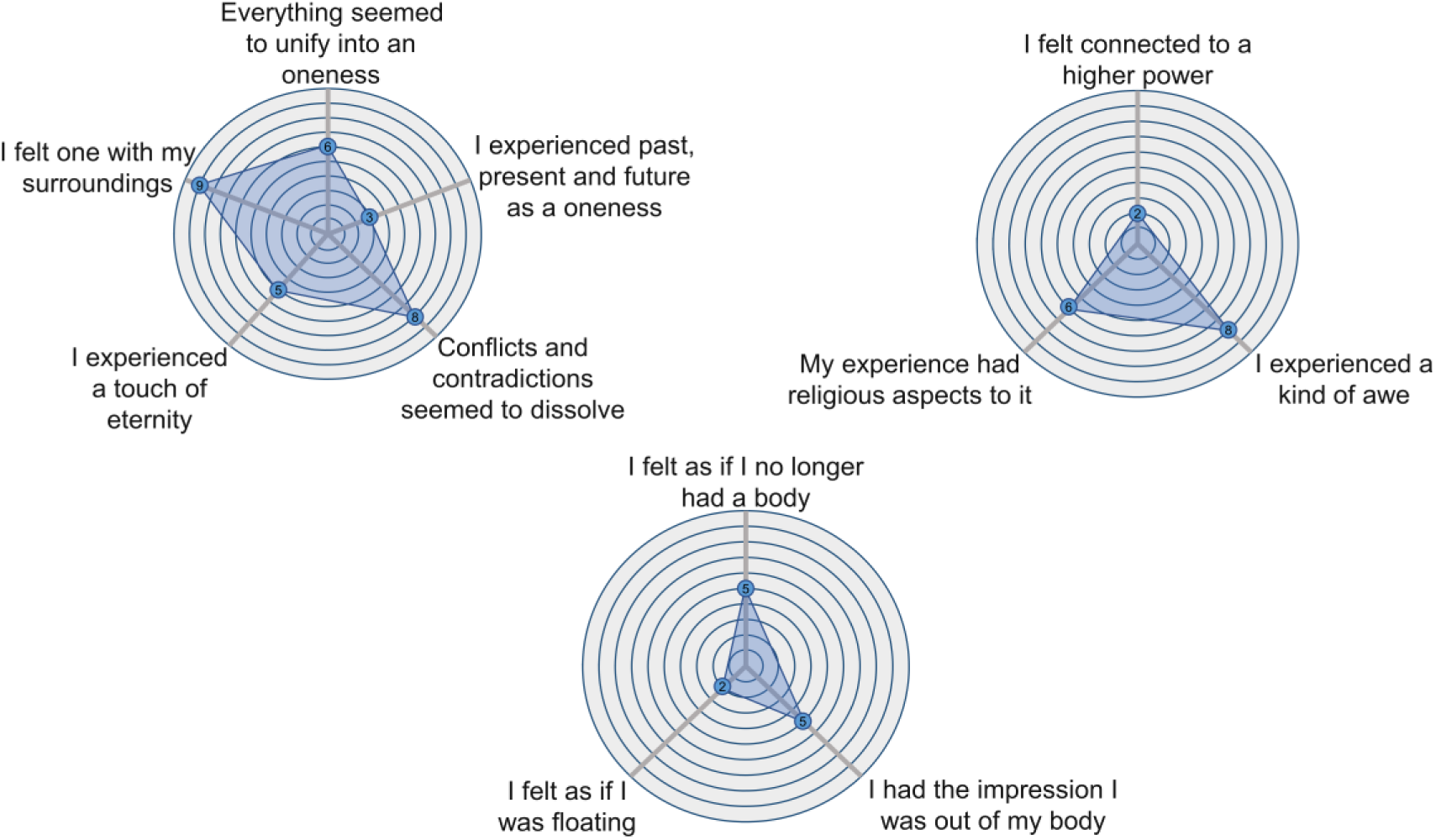
Customizable spider charts used to obtain ratings for the subscales of the 11D-ASC questionnaire. Each spider chart represents a subscale, with as many radii as items. The spider charts were displayed on a touch-sensitive tablet, where participants filled in the subscales by drawing their finger.

### Sound pattern analysis

The sound pattern analysis was conducted by using the Fieldtrip v.20211020 toolbox^[33]^ and Matlab v.2021a. The frequency at which entrainment was expected to be elicited in the recorded EEG signals (i.e., the songs’ beat frequency) was determined in a semi-automatic approach. First, an experienced musician tapped along to each 1-minute song extract on a keyboard, while a custom Matlab program counted the taps and extracted a first approximated beat frequency (i.e., the number of taps per second). The second step involved transforming the songs’ waveforms to the frequency dimension. To this aim, each song was cut from +1 s to +59 s relative to the onset of the musical extract. The first and last seconds of the songs were left out of the analysis to exclude the fade-in and fade-out effects of the songs. The envelopes were extracted using a Hilbert transform, this is, by obtaining the absolute value of the complex-valued analytic signal using the functions ’abs’ and ’hilbert’ in Matlab.

Considering the non-stationarities inherent in our naturalistic music stimulation, the envelopes were transformed into the frequency domain using Welch’s method with a window length of 10 seconds and a 90% overlap, employing a Hanning taper. This transformation was performed using the ’mtmconvolv’ function implemented in Fieldtrip and resulted in a frequency spectrum with a frequency resolution of 0.1 Hz. The exact beat frequency was determined as the spectral amplitude’s peak within a range from -0.5 to +0.5 Hz from the first approximated beat frequency. To explore the temporal dynamics of neural entrainment, the exact beat frequency was measured across time in a separate analysis. The 58-second-long waveforms were segmented into 10 s windows with a 90% overlap and subsequently transformed into the frequency domain with a Hanning taper, as implemented in Fieldtrip’s function ‘mtmconvolv’, yielding frequency spectrums with a frequency resolution of 0.1 Hz. The exact beat frequency for each window was determined as the frequency bin with the maximum spectral amplitude within a range from -0.5 to +0.5 Hz from the approximated beat frequency.

Differences in the spectral amplitude of the beat frequencies between songs were expected and indeed confirmed after the frequency transformations (Fig. 3b). These differences might be explained by there being a different number of beats for each musical extract, which might affect the measures of neural entrainment in an unknown way. To account for these differences, we implemented a normalization procedure. Global entrainment and entrainment across time were measured as a multi-harmonic signal-to-noise ratio (SNR) of the spectral amplitudes of the EEG responses at the beat and significant harmonic frequencies (see section 2.6. EEG data processing and analysis). To normalize these measures, we quantified for each song the multi-harmonic SNR of the spectral amplitudes of the whole waveforms at the beat and significant harmonic frequencies. Subsequently, we computed the ratio between the SNR of the EEG responses and the SNR of the corresponding waveforms. To compute the multi-harmonic SNR of the waveforms, we first summed the spectral amplitudes at the beat frequency (F0), first harmonic (2F0), and second harmonic (3F0) frequency bins (i.e., signal) and at the corresponding neighbor frequencies with one frequency bin of spacing at each flank (i.e., baseline). The multi-harmonic SNR of the waveforms were computed by dividing the signal’s amplitude by the baseline’s amplitude.

### EEG recordings

EEG was continuously recorded during the whole duration of the trials and digitalized at a sampling rate of 1 kHz by a Neuroscan SynAmps RT amplifier (Neuroscan, Compumedics, Charlotte, NC, USA). For the EEG acquisition, 36 sintered electrodes mounted in a neoprene cap (Quick-Cap Neo-Net; Compumedics, Charlotte, NC, USA) at standard locations according to the extended 10-10 system were used. Electrooculograms (EOG) were measured with two bipolar in-cap electrodes placed above and below the right eye (VEOG), and two horizontal in-cap electrodes placed on the outer canthi of the eyes (HEOG). The ground electrode was located at AFz and the common reference electrode between Cz and Cpz. All impedances were kept below 10 kΩ during the whole recording session.

### EEG data processing and analysis

The data analysis was performed offline using the EEGlab v.2021.1 toolbox^[34]^ and the Fieldtrip v.20211020 toolbox^[33]^ running under Matlab v.2021a. For each participant, the continuous recordings were filtered from 0.5 Hz to 45 Hz with a Finite Impulse Response bandpass Kaiser filter to remove slow drifts and line noise, respectively. Excluding the bipolar montages, the filtered data were re-referenced to the average activity of all electrodes to remove any noise from the reference electrode. For each trial, the continuous recordings were segmented from **-**10 s to +60 s relative to the onset of the auditory stimuli. All epochs were merged to detect muscle artifacts based on a semi-automatic approach. First, all epochs were visually inspected with respect to their waveform morphology. Those epochs that included muscle artifacts^[35]^, characterized by high-frequency activity (> 20 Hz) and typically produced by muscular activity near the head, such as swallowing or moving the head, were rejected, unless the artifacts were due to eye movements. Then, independent component analysis (ICA) was computed to remove artifacts produced by eye blinks, eye movements, and heart activity from the EEG signal using the runica algorithm^[36,37]^. Deliberately removing activity in the EEG coming from the heart was critical because the tempo of the songs at 1.65 Hz partly matched the frequency of the human heart, ranging from 1 Hz to 1.67 Hz. Lastly, epochs were baseline corrected by using the EEG activity over the 10 s windows prior to the onset of the songs. The electrodes relative to the vertical and horizontal EOGs were not further included in the analyses.

For each participant and trial, epochs lasting 58 s were sorted by segmenting the pre-processed recordings from +1 to +59 s relative to the onset of the auditory stimuli. Following the same procedure as in a previous study^[38]^, the first second of each epoch was removed: (1) to discard the transient auditory evoked potentials related to the onset of the stimulus^[39–41]^; (2) because steady-state evoke potentials require several cycles of stimulation to be steadily entrained^[42]^; and (3) because several repetitions of the beat are required to elicit a steady perception of beat^[43]^. Additionally, by removing the first and last seconds of the recording, the fade-in and fade-out effects of the stimuli were excluded.

From this step onwards, data were analyzed in two independent strategies. First, to explore *global* measures of entrainment, the obtained 58-second-long epochs were transformed in the frequency domain by using Welch’s method, computed over 10 s windows with a 90% overlap with a Hanning taper, as implemented in Fieldtrip’s function ‘mtmconvolv’ (Fig. 3). This procedure yielded frequency spectrums with a 0.1 Hz frequency resolution. For each participant, song and electrode, the resulting frequency spectrums were averaged across trials. Global entrainment to each song was measured as a multi-harmonic SNR response, meaning that the measure of entrainment was not only limited to the spectral amplitude at the beat frequency. Rather, entrainment was assessed as a multi-harmonic response due to the non-sinusoidal beats of the songs and the nonlinear nature of the brain processes in response to acoustic onsets that might project the neural responses to the beat frequency onto higher harmonics of the beat frequency^[44,45]^.

To determine which harmonics to consider, the significance of each harmonic’s spectral amplitude averaged across participants and electrodes was tested for each song. Specifically, we z-scored the group-level spectral amplitude at the frequency of each harmonic (i.e., signal), with a baseline defined as the corresponding neighboring bins with one frequency bin of spacing, using the formula z(signal) = (signal − baseline mean)/baseline SD. This testing process was carried out sequentially for each harmonic until one harmonic did not reach significance^[46]^. Using this test, the first and second harmonics were considered significant for songs 5 and 6, as they had z-scores > 1.64 (i.e., *p* < 0.05, one-sample, one-tailed test; testing signal > noise). To prevent introducing bias into the results, the same harmonics were employed for computing entrainment across all songs, namely the first and second harmonics. For each participant, song, and electrode we summed the spectral amplitudes at the beat, first harmonic, and second harmonic frequency bins (i.e., signal) and at the baseline bins^[44]^. Lastly, we computed the SNR between the signal and averaged baseline spectral amplitudes.

Considering previous findings indicating that the perception of beat is region-specific^[22,23]^, we examined the scalp-wide distribution of entrainment to identify and select electrodes that are pertinent to the observed entrainment patterns. To this aim, for each song and electrode we averaged the SNR measures across participants and plotted the scalp distribution (Fig. 3d). Upon a visual examination, a consistent fronto-central pattern of entrainment across songs was observed. Therefore, for each participant and song, we quantified global entrainment as a multi-harmonic SNR averaged across fronto-central channels (i.e., FC3, FCz, FC4, C3, Cz, and C4 according to the extended 10-10 standard system). These measures were subsequently normalized by the spectral amplitudes of the songs (see section 2.4. Sound pattern analysis). Lastly, to mitigate potential confound effects related to the idiosyncratic spectral characteristics of the songs, global entrainment was averaged for each participant between songs within tempos.

The second analytical strategy was related to exploring the temporal dynamics of neural entrainment for each participant and song. To this aim, the 58-second-long average epochs were split into 10 s windows with a 90% overlap and subsequently transformed into the frequency domain with a Hanning taper, as implemented in Fieldtrip’s function ‘mtmconvolv’. For each participant, song, electrode and window, the resulting frequency spectrums were averaged across trials, yielding frequency spectrums with a frequency resolution of 0.1 Hz. The deliberate decision of computing FFTs given windows of 10 s are justified by previous literature on entrainment using similar short-lasting epochs^[47,48]^. Also, using overlapping sliding time windows is justified by the fact that we are conducting a fine-grained analysis of entrainment at specific frequencies^[49]^. The significant harmonics for computing entrainment as a multi-harmonic SNR response across time were determined to be the same as in the measurement of global entrainment, namely the first and second harmonics. For each participant, song, electrode and window we summed the spectral amplitudes at the beat, first harmonic, and second harmonic frequency bins (i.e., signal) and baseline bins^[44]^ and computed the SNR. Subsequently, entrainment across time was quantified as the multi-harmonic SNR averaged across fronto-central channels (i.e., FC3, FCz, FC4, C3, Cz, and C4 according to the extended 10-10 standard system). These measures were normalized by the spectral amplitudes of the songs (see section 2.4. Sound pattern analysis). Lastly, entrainment was averaged for each participant and window between songs within tempos.

### Analysis of behavioral measures

#### Reaction time task

Reaction time (RT) was measured as the time in seconds that participants took to respond to the offset of the songs. For each participant, RT was averaged across trials within songs to control for random variability in their performance. For each tempo, RT was averaged between songs to control for possible confound effects provoked by the idiosyncratic musical characteristics of the songs.

#### Go/no-go task

Participants’ responses in the go/no-go task were classified in four types: (a) hits (HTs), if participants pressed the mouse when a go-stimulus was presented; (b) misses (MSs), if participants did not press the mouse when a go-stimulus was presented; (c) correct rejections (CRs), if participants did not press the mouse when a no-go stimulus was presented; and (d) false alarms (FAs), if participants pressed the mouse when a no-go stimulus was presented. For each participant and trial, the proportion of each type of response was calculated by dividing the total number of those responses by the total number of trials within the go/no-go task. The proportions for each type of response were averaged across trials within the same song, and the mean punctuation for each type of response was obtained by calculating the mean across songs within each tempo.

#### 11D-ASC

For each participant, trial and subscale (i.e., Disembodiment, Spiritual experience, and Experience of unity), the average score was computed across items. For each participant, the punctuations in each subscale were averaged across trials within songs. For each tempo, the scores in each subscale were averaged between songs.

## Results

### Reaction time

To explore differences in RT between tempos, a non-parametric Friedman test of differences among repeated measures was conducted. The results show no significant differences in RT between tempos (χ^2^ _(2)_ = 2, *p* = 0.368; M_1.65_ = 1.044, M_2.25_ = 1.208, M_2.85_ = 1.018).

### Go/no-go task

For each of the studied response types (HT, MS, CR and FA), and for each musical tempo, we performed a one-way non-parametric Friedman test of differences among repeated measures. The results show no significant differences between tempos in any response type (HT: χ^2^ _(2)_ = 3.361, *p* = 0.186; M_1.65_ = 0.723, M_2.25_ = 0.728, M_2.85_ = 0.725; MS: χ^2^ _(2)_ = 3.041, *p* = 0.219; M_1.65_ = 0.023, M_2.25_ = 0.019, M_2.85_ = 0.022; CR: χ^2^ _(2)_ = 2.842, *p* = 0.241; M_1.65_ = 0.206, M_2.25_ = 0.202, M_2.85_ = 0.2; FA: χ^2^ _(2)_ = 2.842, *p* = 0.241; M_1.65_ = 0.048, M_2.25_ = 0.051, M_2.85_ = 0.053).

## 11D-ASC

For each subscale of the 11D-ASC (Disembodiment, Spiritual experience, and Experience of unity) and tempo, we conducted a one-way repeated measures Analysis of Variance to investigate differences in these scores between tempos. The results show a significant effect of tempo in Experience of unity (*F*_(2, 36)_ = 4.775, *p* = 0.014, *η^2^* = 0.016). A post-hoc analysis using t-tests revealed a significant difference (*t*_(18)_ = -2.567, *p* = 0.019) between the tempos 1.65 Hz (M_1.65_ = 4.316) and 2.85 Hz (M_2.85_ = 3.646; Fig. 4). No differences were found in the scores of Disembodiment (*F*_(2, 36)_ = 1.309, *p* = 0.277, *η^2^* = 0.000716; M_1.65_ = 3.893, M_2.25_ = 3.444, M_2.85_ = 3.447) and Spiritual Experience (*F*_(2, 36)_ = 0.214, *p* = 0.808, *η^2^* = 0.008; M_1.65_ = 2.579, M_2.25_ = 2.62, M_2.85_ = 2.687).

**Figure 4.**
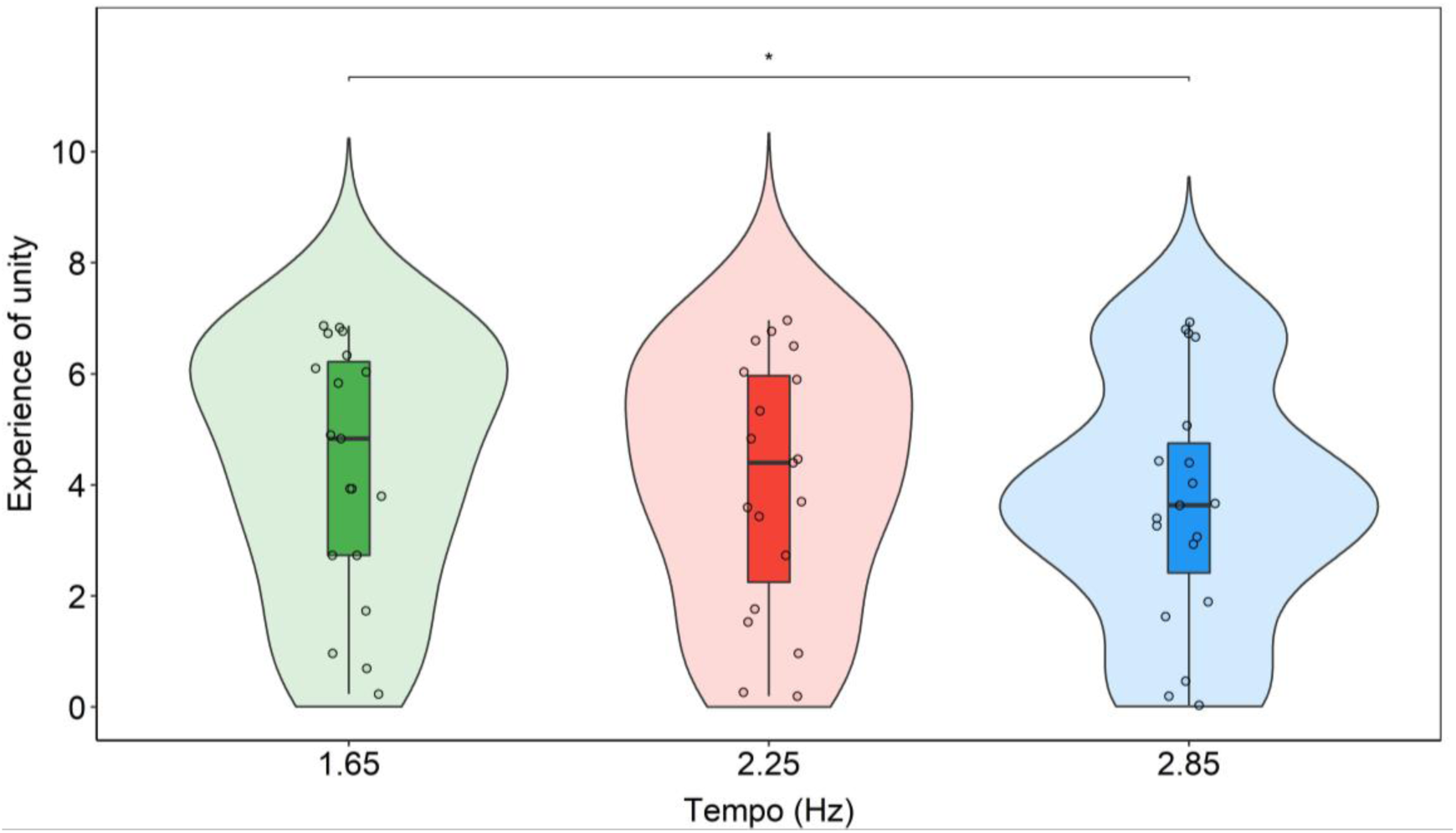
Violin plots showing participants’ scores in the subscale Experience of unity for each tempo condition. Individual data points for each condition are displayed as circular markers. * indicates a p-value lower than 0.05.

### EEG

#### Global entrainment

A linear mixed model analysis was conducted by using the ’lmer’ function from the lme4 package^[50]^ in Rstudio v.4.1.3 to explore the impact of tempo on global entrainment. The covariates "musical training" (indicating whether participants had any form of musical training or not) and "years of musical training" (representing the duration of musical training in years), as assessed with a musical questionnaire, were included in the analysis to control for any potential influence of participants’ musical expertise on entrainment. The normality and dispersion of the residuals were checked. The model included the fixed effects of tempo (with three levels: 1.65, 2.25 and 2.85 Hz), musical training (with two levels: yes or no), and years of musical training. Each participant’s identification code was added as a random effect to account for the correlation among the repeated measurement from the same individual. Adding the covariates improved the fit of the model, as evidenced by a significant decrease of the Bayesian information criterion (BIC) in the model with cowvariates (*BIC* = -84.391) compared to the model without covariates (*BIC* = -80.916; χ^2^_(4)_ = 19.648; *p* < 0.01). The results of the linear mixed model with the main effect of tempo and the covariates about musical training are shown in Table 1.

**Table 1.** Results of the mixed effect model including the main effect of tempo, musical training and years of musical training. Participant identification codes were added as a random effect. Coefficient comparisons for main effects are given as entrainment to 1.65 Hz vs. 2.25 Hz and entrainment to 1.65 Hz vs. 2.85 Hz. ∗ indicates significance level.

| | Coef $\beta$ | SE | t | 95% CI | p |
| --- | --- | --- | --- | --- | --- |
| Intercept | 1.052 | 0.023 | 44.865 | [1.008 1.096] | < 0.001 *** |
| Tempo (1.65 Hz – 2.25 Hz) | -0.065 | 0.030 | -2.182 | [-0.123 -0.007] | 0.036 * |
| Tempo (1.65 Hz – 2.85 Hz) | -0.106 | 0.030 | -3.546 | [-0.164 -0.048] | 0.001 ** |
| Musical Training | -0.040 | 0.040 | -1.007 | [-0.116 0.036] | 0.329 |
| Years of Musical Training | 0.0128 | 0.005 | 2.644 | [0.003 0.022] | 0.018 ** |

To determine the significant comparisons of entrainment between tempos, Tukey’s post-hoc tests with the Kenward-Roger degrees of freedom method were conducted by using the ’emmeans’ function^[51]^ in Rstudio v.4.1.3. Post-hoc contrasts revealed that entrainment was significantly greater for 1.65 Hz compared to 2.85 Hz (*M*_1.65_ = 1.067; *M*_2.85_ = 0.990; *t_(36)_* = 3.546; *p* = 0.003; Fig. 5). No other differences between conditions were found.

**Figure 5.**
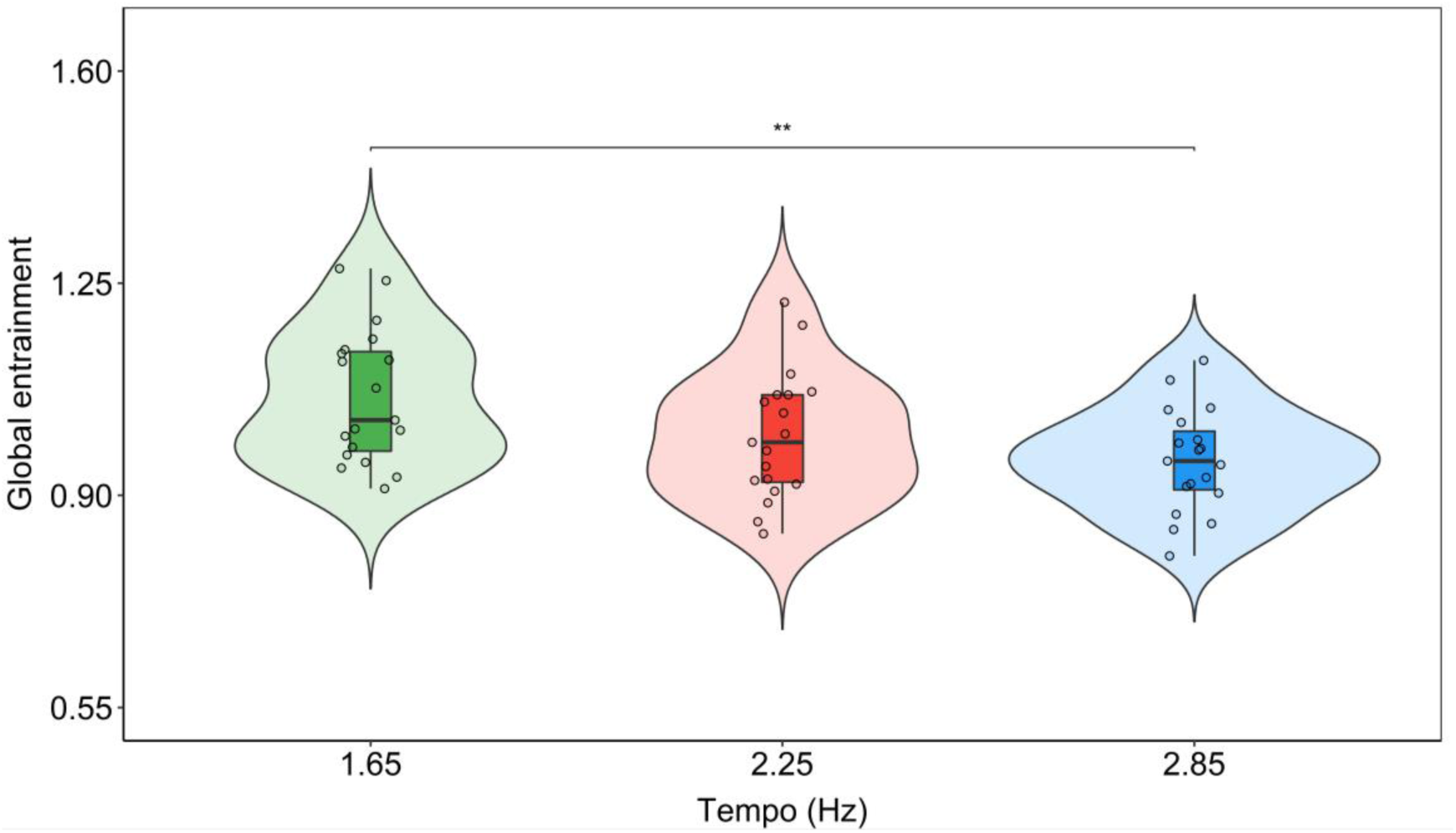
Violin plots showing entrainment to each tempo condition. Entrainment is shown as a multi-harmonic signal-to-noise ratio response normalized by the spectral variations in the musical extracts. Individual data points for each condition are displayed as circular markers. ** indicates a p-value lower than 0.01.

#### Entrainment across time

In order to assess the differences in the temporal dynamics of entrainment to the beat of songs at the different tempos48 one-way repeated measures ANOVAs were conducted, one for each sliding window, with one factor (tempo) and three levels (1.65, 2.25 and 2.85 Hz). A significance level of *p* = 0.01 was implemented to account for the multiple ANOVAs and mitigate the risk of type I error. Depending on the window, post-hoc analyses were conducted using t-tests for normally distributed data or Wilcoxon signed-rank tests for non-normally distributed data. Bonferroni corrections were applied to account for multiple comparisons. The statistical results beyond the main effects are not reported due to the significant quantity of statistical analysis conducted. However, the last time point in which entrainment was measured, which served as the basis for correlational analyses between entrainment and RT and executive function, require further examination. Specifically, a significant effect of tempo was found on the last time point (*χ^2^_(2)_* = 9.789*, p* = 0.007). Post-hoc comparison tests revealed that entrainment was higher for songs at 1.65 Hz compared to both 2.25 Hz (*t_(2)_* = 163*, p* = 0.010) and 2.85 Hz (*t_(2)_* = 165*, p* = 0.010). Fig. 6 visually depicts the results of the main effects across all time points.

**Figure 6.**
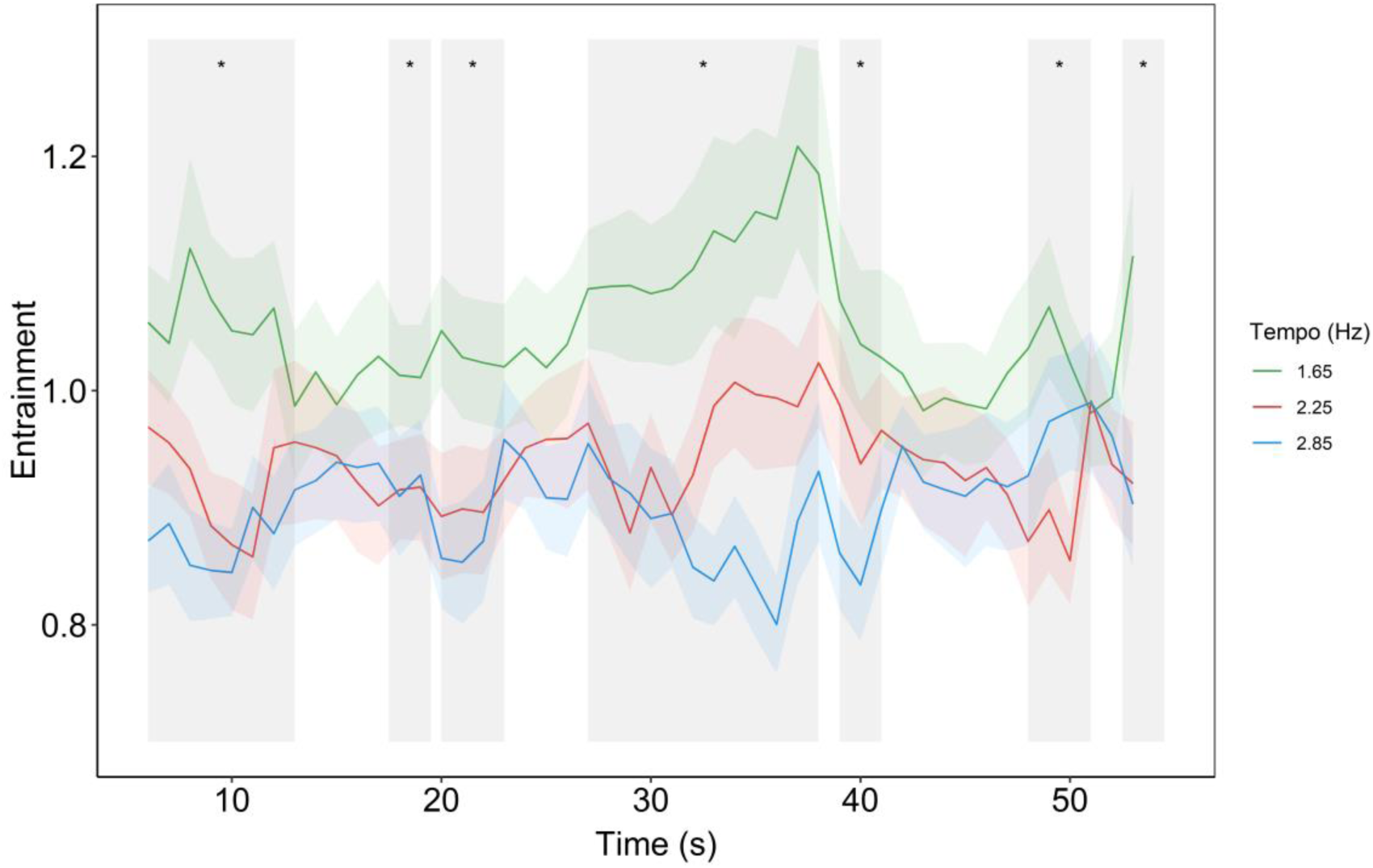
Time-series of entrainment to each tempo condition. Entrainment is shown as a multi-harmonic signal-to-noise ratio response normalized by the spectral variations in the musical extracts. Gray shaded windows represent significant main effects of tempo. * indicates p-value lower than 0.01.

### Brain-behavior correlation analyses

To explore whether the magnitude of entrainment is related to participants’ performance in the objective behavioral measures, linear models were applied to the differences in RT, executive function and entrainment between each pair of tempos. Computing the correlations across differences aimed to uncover whether states in which the brain is highly synchronized to the beats of the songs vs. less synchronized is related to the variability in the participant’s performance between the two conditions. The significance of the relationships was assessed by testing the significance of the slopes of the linear models. To this aim, only entrainment calculated from the last 10-second-long windows were used. This is because participants’ performance in cognitive tasks can be expected to be affected by the brain configuration at the closest moment in time to the task. Also, as the percentage of CR and HT, and the percentage of MS and FS are complementary, correlations with entrainment were only computed for correct rejection and miss responses. The normality of the residuals for each linear model was assessed using Shapiro-Wilk’s tests, revealing that all residuals exhibited a normal distribution. The analysis revealed a significant relationship between the differences in RT and entrainment between tempos 1.65 Hz and 2.25 Hz (*B* = 0.942, *t_(17)_* = 2.276, *p* = 0.027, *R^2^ =* 0.086; Fig. 7a). Also, there was a significant relationship between the differences in CR and entrainment between tempos 1.65 Hz and 2.25 Hz (*B* = -0.018, *t_(17)_* = -2.196, *p* = 0.032, *R^2^ =* 0.081; Fig. 7b) and between 1.65 Hz and 2.85 Hz (*B* = -0.010, *t_(17)_* = -2.118, *p* = 0.039, *R^2^ =* 0.075; Fig. 7c). No other significant relationships were observed.

**Figure 7.**
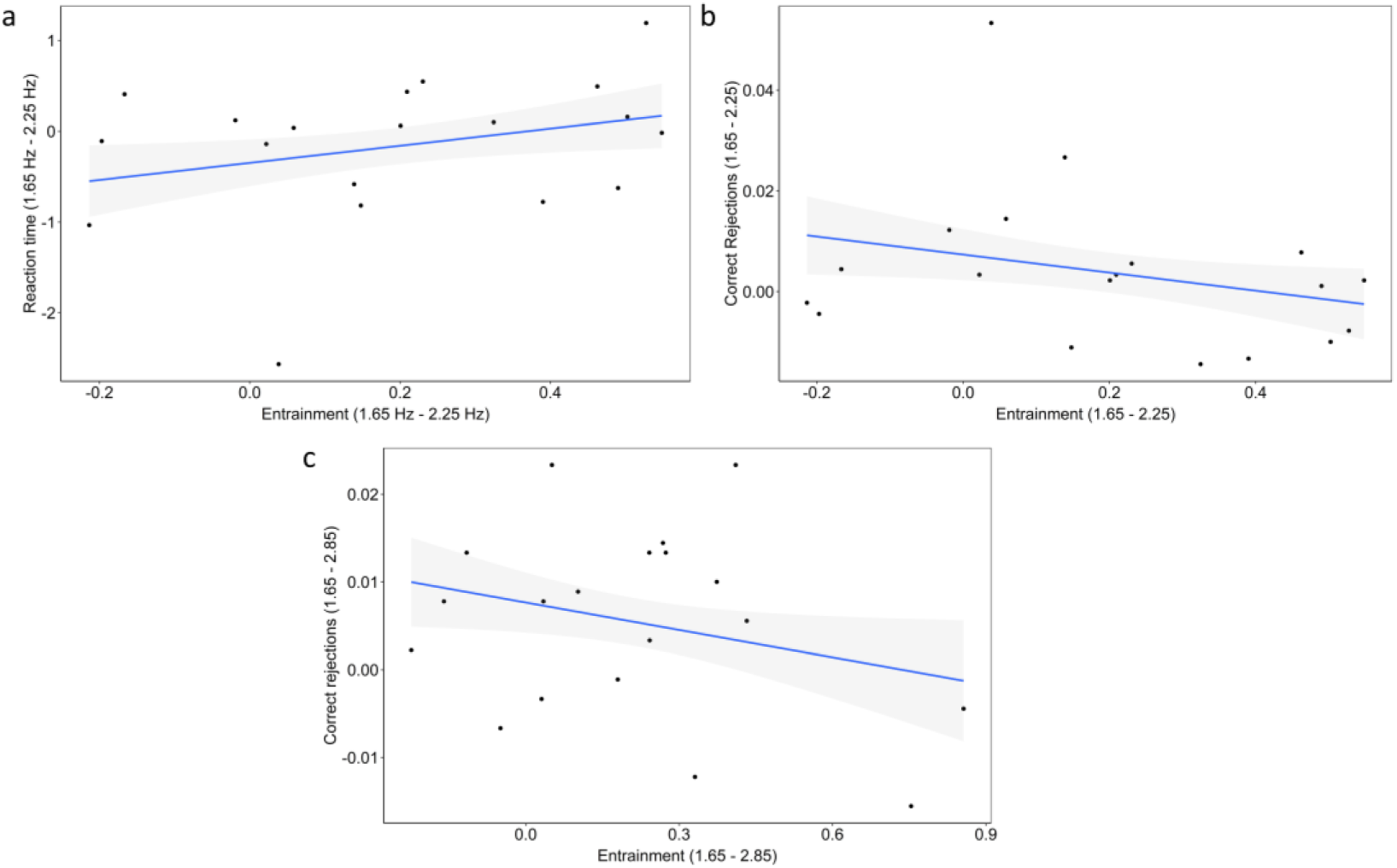
Brain-behavior significant correlations. Relationship between (**a**) the difference in RT and the difference in entrainment between tempos 1.65 Hz and 2.25 Hz, (**b**) the difference in CR and the difference in entrainment between tempos 1.65 Hz and 2.25 Hz, and (**c**) the difference in CR and the difference in entrainment between tempos 1.65 Hz and 2.85 Hz. The regression lines representing the relationships are represented by solid lines, while the shaded areas depict the confidence ranges for each regression at their respective tempos.

To investigate whether the magnitude of entrainment is associated with the subjective experience of participants while listening to the songs, linear models were applied to the differences in participants’ scores in the 11D-ASC subscales and the differences in entrainment between each pair of tempos. In contrast to the objective measures, participant-reported measures of their experience during the stimulation can be expected to be influenced by the overall level of entrainment. No significant relationships were found.

## Discussion

The present study has revealed, using electroencephalography, a relationship between the magnitude of entrainment to the beat of electronic music and some aspects of cognition. In particular, we have shown that the strength of entrainment to the beat of 1-minute-long electronic music can be modulated by the tempo of the music. Our results show that entrainment is higher for stimulation rates at 1.65 Hz compared to faster rates of stimulation, namely 2.85 Hz, but not in comparison to 2.25 Hz. In examining the temporal dynamics of entrainment a similar pattern of results is observed throughout the time course of the songs. Notably, towards the end of the songs, entrainment to the stimulation at 1.65 Hz was higher compared to the other two beat frequencies. The observed neural differences between conditions allowed us to explore whether the magnitude of entrainment is related to differences in cognitive processes or subjective experiences related to altered states of consciousness (ASCs). Although the participants’ reaction time, executive function, feelings of disembodiment, and spiritual experiences were not different depending on the tempo of the music they were exposed to, it was found that music at 1.65 Hz aroused more feelings of unity compared to music at 2.85 Hz. We did not observe any significant relationship between the magnitude of entrainment and participants’ phenomenological experiences during listening to the songs. However, our findings revealed significant yet weak relationships between the magnitude of entrainment and both participants’ reaction time and response inhibition.

Previous empirical evidence showed that entrainment is maximum for periodic auditory stimuli at a rate of 2 Hz within a range of frequencies from 1 to 10 Hz in steps of 1 Hz^[29]^. Our results align with and build upon these findings, having employed stimulus frequencies within a range from 1 to 3 Hz. Specifically, we found more entrainment for music with a beat frequency closer to 2 Hz, namely at 1.65 Hz, compared to music closer to 3 Hz, this is, at 2.85 Hz. We observed that this pattern remains consistent when considering the temporal dynamics of entrainment over the time course of the 60-s musical excerpts. Nevertheless, it is important to consider the large fluctuations in entrainment across time (Fig. 6). These fluctuations could be potentially attributed to stimulus properties, namely variations in the beat accent in the electronic music. However, we contend against this notion due to two main reasons. Firstly, the presence of the beats within the excerpts are quite consistent across time, as visually depicted in Fig. 3a. Secondly, two songs were added for each tempo to lower the effect of the idiosyncratic musical characteristics of each song on the measures of entrainment, attenuating the influence of beat-related stimulus properties. It remains unclear why entrainment to beat frequencies of 2.25 Hz, which is closer to 2 Hz compared to 1.65 Hz, do not exhibit the highest level of entrainment across the tempos used in the study. The predominantly fronto-central scalp distribution of entrainment also aligns with previous findings^[52]^. This pattern can be attributed to the proximity of fronto-central channels to neural generators involved in sensorimotor processing, such as the sensorimotor cortex, and the supplementary motor area, as well as the auditory system.

Given that entrainment is modulated by the tempo of music, we suggest that complex brain mechanisms might be tuning entrainment, most probably in favor of brain function^[19,20]^. The stimulation frequency of 1.65 Hz partially matches the human optimal rate for sensorimotor behavior^[53–56]^. Previous research discussing this match has proposed the existence of a brain mechanism facilitating auditory-motor behavior through entrainment when being presented with auditory stimuli at rates around 2 Hz^[29]^ that could also explain part of our results. However, the likelihood of small unintentional body movements while participants were listening to the music cannot be discounted. Body movement may account for the differences in entrainment between tempos, as such movement might be amplified for the songs at the tempo closer to 2 Hz. Upcoming studies should monitor small body movements, especially head movement, during music listening when exploring entrainment.

To the best of our knowledge, no previous studies had explored how the rate of repetitive auditory stimuli might modulate ASCs’ characteristics. Here, we measured proxies of ASCs both in terms of cognitive function (i.e., reaction time and executive function) and in terms of subjective experience (i.e., by using three subscales of the 11D-ASC)^[28]^ while listening to the music at different tempos. We found that the tempo of electronic music did not affect participants’ overall reaction time or executive capacities. Similarly, participants’ experience of unity and feelings of disembodiment did not change depending on the tempo of the music. These results suggest that the presentation rate of repetitive stimuli does not affect differently these aspects of cognitive function and human experience. Noteworthy, these results do not indicate whether cognition and human experience are altered under the conditions participants where in, as only the rate of stimulation is being accounted for. However, participants felt more experiences of unity for the music at 1.65 Hz compared to the music at 2.85 Hz, mirroring the entrainment findings and suggesting a potential brain-behavior relationship. One limitation of the study is the potential impact of the course of the experimental procedure on the participants’ phenomenological ratings. Specifically, possible confound effects on the phenomenological experiences reported might rise from having participants conducting the go/no-go task in between listening to the music extracts and filling out the retrospective 11D-ASC questionnaire.

In previous literature on ASCs, the usage of repetitive stimulation to trigger altered mental states^[7–9,14,15]^ has been explained by entrainment^[17]^, but with no direct evidence to support that claim.

Critically, for the first time this study has explored a relationship between the magnitude of entrainment and ASCs. We found three weak, yet significant, brain-behavior associations. Given the modest strength of these relationships, caution should be exercised in interpreting the results and further investigation is warranted to elucidate potential underlying mechanisms. To explore these associations, the neural metric employed was entrainment observed during the final 10 seconds of stimulation, which was higher for songs at 1.65 Hz compared to both songs at 2.25 Hz and 2.85 Hz. First, we found that the more differences in entrainment between songs at 1.65 Hz and 2.25 Hz, the more differences in participants’ reaction time to the offset of these songs. In our study, reaction time was used as a measure related to the level of absorption to the music, indicating participants’ cognitive responsiveness and engagement with the auditory stimuli. Because rhythmically-induced ASCs are characterized by a selective focus on environmental stimuli^[11]^, reaction time to the offset of the songs is a potential proxy of ASCs. While entrainment was higher for songs at 1.65 Hz compared to songs at 2.25 Hz, the differences in reaction time between these tempos did not exhibit a consistent trend among participants. This is evidenced by the null differences in reaction time across tempos. Also, Fig. 7 shows that differences in reaction time between songs at 1.65 Hz and songs at 2.25 Hz are distributed above and below zero. In other words, our results show that higher levels of neural synchronization are related to both faster and slower reaction times, depending on the participant. These findings invite further investigation into the relationship among entrainment, reaction times, and personality traits and/or individual cognitive characteristics. One such significant trait could be musical training, as it was significant in explaining a portion of the observed variance in the strength of entrainment across conditions. However, the limited representation of participants with musical training within our sample precluded a comprehensive investigation into its potential influence on the observed associations.

Additional brain-behavior associations revealed that the more differences in entrainment participants showed between tempos 1.65 Hz and 2.25 Hz, the more similar participants’ inhibition responses between the two conditions were. The same pattern was found between tempos 1.65 Hz and 2.85 Hz. The inhibition responses were measured as participants’ correct rejections in the go/no-go task performed after listening to the musical extracts. Empirical evidence suggests that the inhibition response relies on neural activity involving the pre-supplementary motor area^[57]^. Also, previous research has shown persistent effects of oscillatory entrainment on cognition^[58]^. In our study, the fronto-central scalp distribution of entrainment observed during listening to the songs suggest that motor areas, including the supplementary motor area, might be recruited. Therefore, it is a possibility that a higher strength of entrainment in motor areas during listening to the songs might decrease neural variability during correct rejection responses in the go/no-go task, resulting in fewer differences in participants’ inhibition response. While this holds physiological interest, its significance in relation to the hypothesized correlation between entrainment and ASCs appears limited. Nevertheless, it does imply that repetitive stimulation effectively entrains regions associated with inhibitory responses, potentially influencing related behavioral outcomes. Consequently, future investigations should delve into the potential correlation between entrainment, inhibitory responses, and ASCs.

## Conclusions

To our knowledge, this is the first report of a relationship between entrainment and ASCs, although this relationship has been suggested in previous literature. In summary, this article has argued that entrainment and the phenomenological aspects of ASCs induced by repetitive stimuli are related. Our results showing that entrainment is higher for stimulation rates at 1.65 Hz are broadly consistent with previous findings. We found an association between entrainment and absorption to the music, as measured with participants’ reaction times to the offset of the stimulation. Specifically, we observed that the more the strength of participants’ entrainment to the music, as measured by differences in entrainment between songs at 1.65 Hz and 2.25 Hz, the more differences in participants’ reaction time to the offset of these songs. While entrainment was higher for songs at 1.65 Hz compared to 2.25 Hz, reaction time was the same across tempos. Therefore, we suggest that individual personality or cognitive traits might be modulating whether the strength of entrainment is related to more or less time to respond and, subsequently, to whether participants are more or less absorbed by the songs. Although musical training emerged as a significant factor explaining variance in the magnitude of entrainment, the small subset of participants with musical training in our sample limits deeper exploration of its impact on this brain-behavior association. Additionally, given the weak strength of this relationship, caution is advised when interpreting these findings.

## Data availability

The data and analysis code will be publicly available here upon publication (https://osf.io/nkzcg/).

## Authors contributions

**RAT**: Conceptualization, Methodology, Software, Formal analysis, Investigation, Data curation, Writing – original draft, Writing – review & editing, Visualization, Supervision, Project administration. **SLM**: Conceptualization, Methodology, Writing – review & editing. **MDA**: Conceptualization, Writing – review & editing, Supervision, Project administration, Funding acquisition. **CE**: Conceptualization, Methodology, Writing – review & editing, Supervision, Project administration, Funding acquisition.

## References

1. St John, G. Liminal being: Electronic dance music cultures, ritualization and the case of psytrance in The Sage Handbook of Popular Music (eds. Bennet, A. & Waksman, S.) 243–260 (Sage, 2015).

2. Tramacchi, D. Field tripping: psychedelic communitas and ritual in the Australian Bush. Journal of Contemporary Religion 15, 201–213 (2000).

3. Newson, M., Khurana, R., Cazorla, F. & van Mulukom, V. ‘I get high with a little help from my friends’ - how raves can invoke identity fusion and lasting co-operation via transformative experiences. Front Psychol 12, (2021).

4. Hood, R. W. J. Methodological issues in the use of psychedelics in religious rituals in Seeking the Sacred With Psychoactive Substances: Chemical Paths to Spirituality and to God: History and Practices; Insights, Arguments and Controversies (ed. Ellens, J. H.) 395–410 (Bloomsbury, 2014).

5. Nencini, P. The shaman and the rave party: social pharmacology of ecstasy. Subst Use Misuse 37, 923–939 (2002).

6. Gingras, B., Pohler, G. & Fitch, W. T. Exploring shamanic journeying: repetitive drumming with shamanic instructions induces specific subjective experiences but no larger cortisol decrease than instrumental meditation music. PLoS One 9, e102103 (2014).

7. Kjellgren, A. & Eriksson, A. Altered states during shamanic drumming: a phenomenological study. International Journal of Transpersonal Studies 29, 1–10 (2010).

8. Szabó, C. The effects of listening to monotonous drumming on subjective experiences in Music and Altered States (eds. Aldridge, D. & Fachner, J.) 51–59 (Kingsley, 2006).

9. Walker, M. Music as knowledge in shamanism and other healing traditions of siberia. Arctic Anthropol 40, 40–48 (2003).

10. Goodman, F. D. Where The Spirits Ride The Wind: Trance Journeys And Other Ecstatic Experiences. (Indiana University Press, 1990).

11. American Psychiatric Association. Diagnostic And Statistical Manual Of Mental Disorders. (American Psychiatric Publishing, 1994).

12. Hutson, S. R. Technoshamanism: spiritual healing in the rave subculture. Popular Music and Society 23, 53–77 (1999).

13. Schäfer, T., Fachner, J. & Smukalla, M. Changes in the representation of space and time while listening to music. Front Psychol 4, 51929 (2013).

14. 15. Harner, M. The Way Of The Shaman. (Harper & Row, 1990).

15. Krippner, S. The epistemology and technologies of shamanic states of consciousness. Journal of Consciousness Studies **11–12**, 93–118 (2000).

16. Winkelman, M. A paradigm for understanding altered consciousness: the integrative mode of consciousness in Altering Consciousness Multidisciplinary Perspectives (eds. Cardeña, E. & Winkelman, M.) 23–41 (Praeger, 2011).

17. Vaitl, D. et al. Psychobiology of altered states of consciousness. Psychol Bull 131, 98–127 (2005).

18. Large, E. W. Resonating to musical rhythm: theory and experiment in Psychology of time (ed. Grondin, S.) 189–232 (Emerald Group, 2008).

19. Bittman, E. L. Entrainment is NOT synchronization: an important distinction and its implications. J Biol Rhythms 36, 196–199 (2021).

20. Lakatos, P., Karmos, G., Mehta, A. D., Ulbert, I. & Schroeder, C. E. Entrainment of neuronal oscillations as a mechanism of attentional selection. Science *(1979)* 320, 110– 113 (2008).

21. Zoefel, B., ten Oever, S. & Sack, A. T. The involvement of endogenous neural oscillations in the processing of rhythmic input: more than a regular repetition of evoked neural responses. Front Neurosci 12, (2018).

22. Chen, J. L., Penhune, V. B. & Zatorre, R. J. Listening to musical rhythms recruits motor regions of the brain. Cerebral Cortex 18, 2844–2854 (2008).

23. Grahn, J. A. & Brett, M. Rhythm and beat perception in motor areas of the brain. J Cogn Neurosci 19, 893–906 (2007).

24. Gosseries, O. et al. Behavioural and brain responses in cognitive trance: A TMS-EEG case study. Clinical Neurophysiology 131, 586–588 (2020).

25. Hove, M. J. et al. Brain network reconfiguration and perceptual decoupling during an absorptive state of consciousness. Cerebral Cortex 26, 3116–3124 (2016).

26. Kawai, N., Honda, M., Nishina, E., Yagi, R. & Oohashi, T. Electroencephalogram characteristics during possession trances in healthy individuals. Neuroreport 28, 949–955 (2017).

27. Oohashi, T. et al. Electroencephalographic measurement of possession trance in the field. Clinical Neurophysiology 113, 435–445 (2002).

28. Studerus, E., Gamma, A. & Vollenweider, F. X. Psychometric evaluation of the altered states of consciousness rating scale (OAV). PLoS One 5, (2010).

29. Will, U. & Berg, E. Brain wave synchronization and entrainment to periodic acoustic stimuli. Neurosci Lett 424, 55–60 (2007).

30. Sevenius Nilsen, A. E., Juel, B. E., Farnes, N., Romundstad, L. & Storm, J. F. Behavioral effects of sub-anesthetic ketamine in a go/no-go task. J Psychedelic Stud 4, 156–162 (2020).

31. Young, M. E., Sutherland, S. C. & McCoy, A. W. Optimal go/no-go ratios to maximize false alarms. Behav Res Methods 50, 1020–1029 (2018).

32. López-Mochales, S., Aparicio-Terrés, R., Díaz-Andreu, M. & Escera, C. Acoustic perception and emotion evocation by rock art soundscapes of Altai (Russia). Front Psychol 14, (2023).

33. Oostenveld, R., Fries, P., Maris, E. & Schoffelen, J. M. FieldTrip: Open source software for advanced analysis of MEG, EEG, and invasive electrophysiological data. Comput Intell Neurosci 2011, (2011).

34. Delorme, A. & Makeig, S. EEGLAB: an open source toolbox for analysis of single-trial EEG dynamics including independent component analysis. J Neurosci Methods 134, 9–21 (2004).

35. Goncharova, I. I., McFarland, D. J., Vaughan, T. M. & Wolpaw, J. R. EMG contamination of EEG: spectral and topographical characteristics. Clinical Neurophysiology 114, 1580– 1593 (2003).

36. Bell, A. J. & Sejnowski, T. J. An information-maximization approach to blind separation and blind deconvolution. Neural Comput 7, 1129–1159 (1995).

37. Makeig, S., Jung, T. P., Bell, A. J., Ghahremani, D. & Sejnowski, T. J. Blind separation of auditory event-related brain responses into independent components. Proc Natl Acad Sci U S A 94, 10979–10984 (1997).

38. Nozaradan, S., Peretz, I. & Mouraux, A. Selective neuronal entrainment to the beat and meter embedded in a musical rhythm. Journal of Neuroscience 32, 17572–17581 (2012).

39. Saupe, K., Schröger, E., Andersen, S. K. & Müller, M. M. Neural mechanisms of intermodal sustained selective attention with concurrently presented auditory and visual stimuli. Front Hum Neurosci 3, 688 (2009).

40. Nozaradan, S., Peretz, I., Missal, M. & Mouraux, A. Tagging the neuronal entrainment to beat and meter. Journal of Neuroscience 31, 10234–10240 (2011).

41. Nozaradan, S., Peretz, I. & Mouraux, A. Steady-state evoked potentials as an index of multisensory temporal binding. Neuroimage 60, 21–28 (2012).

42. Regan, D. Human brain electrophysiology: evoked potentials and evoked magnetic fields in science and medicine. Elsevier. (1989).

43. Repp, B. H. Sensorimotor synchronization: A review of the tapping literature. Psychon Bull Rev 12, 969–992 (2005).

44. Retter, T. L., Rossion, B. & Schiltz, C. Harmonic amplitude summation for frequency-tagging analysis. J Cogn Neurosci 33, 2372–2393 (2021).

45. Zhou, H., Melloni, L., Poeppel, D. & Ding, N. Interpretations of frequency domain analyses of neural entrainment: periodicity, fundamental frequency, and harmonics. Front Hum Neurosci 10, (2016).

46. Lenc, T., Keller, P. E., Varlet, M. & Nozaradan, S. Neural tracking of the musical beat is enhanced by low-frequency sounds. Proc Natl Acad Sci U S A 115, 8221–8226 (2018).

47. Okawa, H., Suefusa, K. & Tanaka, T. Neural entrainment to auditory imagery of rhythms. Front Hum Neurosci 11, (2017).

48. Tal, I. et al. Neural entrainment to the beat: the “missing-pulse” phenomenon. Journal of Neuroscience 37, 6331–6341 (2017).

49. Batterink, L. J. & Choi, D. Optimizing steady-state responses to index statistical learning: Response to Benjamin and colleagues. Cortex 142, 379–388 (2021).

50. Bates, D., Mächler, M., Bolker, B. M. & Walker, S. C. Fitting linear mixed-effects models using lme4. J Stat Softw 67, (2014).

51. Lenth, R. Emmeans: Estimated marginal means, aka least-squares means. R package version 1.4.2. Preprint at (2019).

52. Tierney, A. & Kraus, N. Neural entrainment to the rhythmic structure of music. J Cogn Neurosci 27, 400–408 (2015).

53. Drake, C., Jones, M. R. & Baruch, C. The development of rhythmic attending in auditory sequences: attunement, referent period, focal attending. Cognition 77, 251–288 (2000).

54. Large, E. W. & Jones, M. R. The dynamics of attending: how people track time-varying events. Psychol Rev 106, 119–159 (1999).

55. McAuley, J. D. & Jones, M. R. Modeling effects of rhythmic context on perceived duration: a comparison of interval and entrainment approaches to short-interval timing. J Exp Psychol Hum Percept Perform 29, 1102–1125 (2003).

56. van Noorden, L. & Moelants, D. Resonance in the perception of musical pulse. Int J Phytoremediation 21, 43–66 (1999).

57. Sánchez-Carmona, A. J., Santaniello, G., Capilla, A., Hinojosa, J. A. & Albert, J. Oscillatory brain mechanisms supporting response cancellation in selective stopping strategies. Neuroimage 197, 295–305 (2019).

58. Roberts, B. M., Clarke, A., Addante, R. J. & Ranganath, C. Entrainment enhances theta oscillations and improves episodic memory. Cogn Neurosci 9, 181–193 (2018).

